# Underlying microevolutionary processes parallel macroevolutionary patterns in ancient Neotropical Mountains

**DOI:** 10.1101/2020.08.26.268870

**Authors:** Marcos Vinicius Dantas-Queiroz, Tami da Costa Cacossi, Bárbara Simões Santos Leal, Cleber Juliano Neves Chaves, Thais N. C. Vasconcelos, Leonardo de Melo Versieux, Clarisse Palma-Silva

## Abstract

**Aim:** The exceptional species-richness associated with mountains worldwide is linked to the fragmented topography of these areas, responsible for constantly isolating populations during periods of climatic fluctuations. Consequently, endemism and spatial turnover in mountains are very high and few species are widespread among entire mountain ranges, precluding population-level studies that help understanding how macroevolutionary patterns were shaped. Here, we used the bromeliad *Vriesea oligantha*, a species endemic to, but widespread in, one of the most species-rich ancient montane areas in the globe, the Espinhaco Range, to test how environmental changes over time may have acted on the evolutionary history of this taxon, contributing to understanding how montane macroevolutionary patterns were shaped. Through analyses of plastidial and nuclear DNA of *V. oligantha*, we dated its origin and intraspecific diversification, and estimated the genetic diversity, structure and migration rates among populations. Using climatic and geographic variables, we modeled suitable areas for the present and the past, estimating corridors between isolated populations. We also used demographic analyses to estimate ancient population dynamics of *V. oligantha*. Finally, we tested whether climatic variables or geographical distance explain the observed population structure. The origin and intraspecific diversification of *V. oligantha* are related to early climatic oscillations during the Plio-Pleistocene. This species has a high population structure due to its low pollen and seed dispersibility. The analysis of species distribution modeling estimated corridors between populations in the past, whereas the structure of *V. oligantha* results from both models of isolation by distance and isolation by environment. The phylogeographic patterns of *Vriesea oligantha* reflect previously recognized spatial and temporal macroevolutionary patterns in the Espinhaco Range, providing insights into how microevolutionary processes may have given rise to this astonishing mountain biodiversity.

## INTRODUCION

Mountains are remarkable models for evolutionary studies since they host a substantial proportion of the world’s biodiversity and harbors high levels of endemism (Antonelli et al., 2018; Perrigo, Hoorn & Antonelli, 2020). The drivers of this diversity began to be explored by Alexander von Humboldt, who studied the relationship between mountain vegetation and its abiotic traits (Humboldt, 1807). But how this astonishing biodiversity arises is still a debatable subject.

Based on the hypothesis of Quaternary refugies (Haffer, 1969; Vanzolini & Williams, 1970), climatic fluctuations are one of the main factors that would explain the high biodiversity in mountains (Antonelli et al., 2018; Rull, 2011). Accordingly, extant species restricted to mountain tops are in interglacial refuges, while in glacial periods, their area of distribution would be larger, seeking their optimal niches towards lowlands (Perrigo et al., 2020). The Last Glacial Maximum (LGM, ~21 kyr) predominates in the literature as a common driver of diversification, mainly on Northern but also in Southern latitudes (Beheregaray, 2008; Feliner, 2011) although repeated cycles of climatic oscillations from the last million years may also have contributed to isolate mountain populations, leading to genetic drift and local adaptation to new environments and potentially generating new species (Flantua, O’Dea, Onstein, Giraldo & Hooghiemstra, 2019). This phenomenon has been described as a species pump (Haffer, 1997), biodiversity pump (Rull, 2005), or isolation-cooling hypothesis (Rull & Vegas-Vilarrúbia, 2020), and is thought to be associated with the elevated speciation rates linked to mountain ranges.

An example of the link between increased diversification during periods of climatic fluctuations can be observed in the Espinhaço Range in Eastern South America. The Espinhaço Range harbors an astonishing plant diversity, accounting for nearly 15% of the entire Brazilian Flora, with ca. 2,000 endemic species, making these mountains home of one of the highest species richness and endemism rates of the world (Silveira, Dayrell, Fiorini, Negreiros & Borba, 2020). Many Espinhaço endemic angiosperms lineages exhibit high rates of diversification in the last 5 Myr (Vasconcelos et al., 2020), despite the ancient age of these mountains (640 Myr - Precambrian) (Pedreira & de Waele, 2008), indicating the species-pump phenomenon caused by the climatic oscillations of the Plio-Pleistocene period as one of the most likely drivers of the Espinhaço biota formation (Alcantara, Ree & Mello-Silva, 2018; Ribeiro, Rapini, Damascena & van Den Berg, 2014).

The species-pump assumes that spatial isolation is one of the main drivers of speciation, and a presumable consequence is that distributions of endemic species are clustered in small areas (Vasconcelos et al., 2020), leading to high spatial turnover. This high degree of endemism associated to different portions of the Espinhaço Range is also remarkably congruent between different organisms (Chaves, Freitas, Vasconcelos & Santos, 2015; Echternacht, Trovó, Oliveira & Pirani, 2011) and has led to the recognition of distinct biogeographical regions in the northern (e.g. the Chapada Diamantina province) and in the mid-southern portions of the range (e.g. the Diamantina Plateau and the Iron Quadrangle districts) (Colli-Silva et al., 2019).

Are past climatic fluctuations common factors that could explain the origin of these congruent biogeographic and diversification patterns? If on a macroevolutionary scale the species-pump boosts diversification rates through population isolation events, one can expect to observe microevolutionary processes (i.e., genetic drift and restricted gene flow) acting at the population level in the early stages of divergence (Li, Huang, Sukumaran & Knowles, 2018). Confirm the effects of such processes in already diverged lineages may be unfeasible, but perhaps it might be possible to observe these first steps of speciation in extant populations of the same species distributed among isolated mountains (Pinheiro et al., 2013). Even though there have been important contributions testing the species-pump hypothesis in other montane areas (e.g., Andes: Sedano & Burns, 2010; Himalayas: Liu et al., 2016; Sierra Nevada: Schoville, Roderick & Kavanaugh, 2012) studies in the Espinhaço Range are often prevented by the narrow distributions of endemic species, precluding the understanding of how underlying microevolutionary processes have promoted the emergence of macroevolutionary patterns of these mountains.

In this context, we used the wide-distributed Espinhaço-endemic bromeliad *Vriesea oligantha* species complex as a model system to investigate how climatic oscillations have shaped evolution of endemic lineages in this mountain range. Following the wide distribution and high-altitude suitability of *V. oligantha*, we hypothesized that its evolutionary history likely follows the history of its habitat, varying in area and connectivity due to ancient climate oscillations. To test this premise, we predict that (i) isolation by distance together with isolation by environment are the main factors driving diversification, in accordance with the species-pump model, and (ii) past corridors connecting populations may have existed when climate was cooler. If both hypotheses are corroborated, we can expect that (iii) the genetic structure of populations reflect the macroevolutionary patterns found in lineages endemic to the Espinhaço Range. To answer these questions, we adopted different approaches, including population genetics, dated phylogenies and ecological niche modeling.

## MATERIAL AND METHODS

### Sampling and DNA extraction

The species complex *Vriesea oligantha* comprises morphologically similar species (the “*limae*” clade (Machado et al., 2019), all rupicolous and epiphytic bromeliads endemic to the Espinhaço Range. For the purpose of this work, we considered all sampled individuals as *Vriesea oligantha,* sampling 229 individuals from 14 populations (Table 1), covering its entire distribution range (Figure 1).

**Figure 1.**
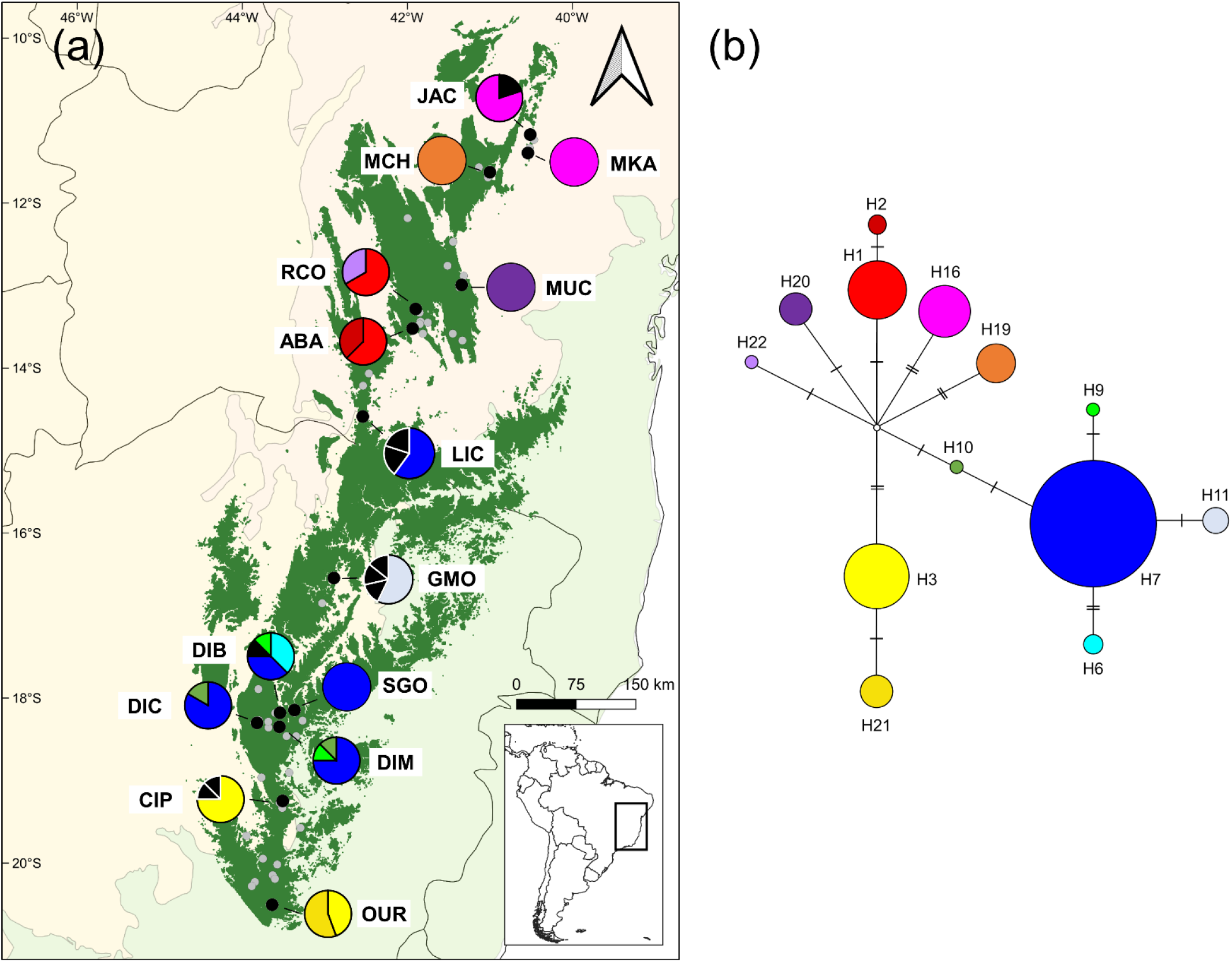
(a) Map of the Espinhaço Range (dark green) and its surrounding phytogeographic domains: Caatinga (North, pink), Cerrado (West, beige) and Atlantic Forest (East and South, light green). Dark lines are the Brazilian States borders. This map shows the entire geographic distribution of *Vriesea oligantha*. Grey points represent occurrence data from herbarium specimens and black points are the sampled populations (See Table 1, for abbreviation codes). The pie charts represent the frequency of occurrence of each haplotype in each population; haplotype colors correspond to those shown in the network; black haplotypes are singletons. (b) Median-joining network depicting the relationships among the haplotypes of *V. oligantha*.

**Table 1.** Population names, geographic locality and tested individual samples for plastidial and nuclear data.

| **Population** | **Location** | **Latitude** | **Longitude** | **Altitude (m)** | **Plastidial *n*** | **Nuclear *n*** |
| --- | --- | --- | --- | --- | --- | --- |
| JAC | Jacobina - BA | -11.1704 | -40.5112 | 713 | 5 | 16 |
| MKA | Miguel Calmon - BA | -11.3933 | -40.5393 | 1003 | 4 | - |
| MCH | Morro do Chapéu - BA | -11.6263 | -41.0000 | 904 | 6 | 19 |
| MUC | Mucugê - BA | -12.9915 | -41.3440 | 980 | 5 | 19 |
| RCO | Rio de Contas - BA | -13.5244 | -41.9567 | 1577 | 6 | 30 |
| ABA | Abaíra - BA | -13.2838 | -41.8998 | 1654 | 8 | 19 |
| LIC | Licínio de Almeida - BA | -14.5874 | -42.5400 | 915 | 5 | 5 |
| GMO | Grão Mogol - MG | -16.5447 | -42.8910 | 1088 | 7 | 20 |
| DIB | Diamantina (Biribiri) - MG | -18.1787 | -43.5447 | 1339 | 8 | 20 |
| DIC | Diamantina (Cons. Mata) - MG | -18.3012 | -43.8224 | 1313 | 6 | 18 |
| DIM | Diamantina (Milho Verde) - MG | -18.3490 | -43.5512 | 1206 | 8 | - |
| SGO | São Gonçalo do Rio Preto - MG | -18.1460 | -43.3687 | 906 | 10 | 15 |
| CIP | Conceição do Mato Dentro - MG | -19.2467 | -43.5116 | 1270 | 8 | 23 |
| OUR | Ouro Branco - MG | -20.5051 | -43.6390 | 1375 | 9 | 25 |

We extracted the total genomic DNA from silica-gel-dried leaves following a modified CTAB protocol described by Tel-Zur, Abbo, Myslabodski & Mizrahi (1999). Initially we tested ten plastidial markers (*petA-psbJ, petG-trnC, psbA-trnH, rpoB-trnC, psbM-trnD, rpoB-trnC-petN, trnK-matK-trnK, trnL-trnF, ycf1, ycf6-trnC*) and one nuclear (*phyC*), selecting two plastidial markers based on the higher polymorphism levels, *ycf1* (Barfuss et al., 2016) and *trnL-trnF* (Barfuss, Samuel, Till & Stuessy, 2005) to be amplified in 95 samples from 14 populations.

### Plastidial DNA sequencing and nuclear microsatellite genotyping

Amplifications for the *ycf1* and *trnL-trnF* markers were performed in a 30 μL reaction using 3 ng of genomic DNA, 5x GoTaq Green Master Mix (Promega, Madison, USA), 0.5 μM of each primer and 1% of DMSO. Polymerase chain reaction (PCR) was conducted using a Veriti 96-well Thermal Cycler (Applied Biosystems, Foster City, USA) following the protocols described in Barfuss et al. (2005, 2016). PCR products were then purified and sequenced both forward and reverse directions on Macrogen (Seoul, South Korea). Consensus sequences and the alignment matrix were assembled on Geneious R10, using the default aligner algorithm. Indels longer than 1 base pair were removed due to uncertain homology. Both markers were concatenated for subsequent analyses. Sequences are deposited in GenBank (XXXXXX-YYYYYY).

For nuclear microsatellites (nrSSR) loci, we genotyped nine polymorphic markers previously transferred from other bromeliads (Cacossi, Dantas-Queiroz & Palma-Silva, 2019) in 229 individuals from 12 populations. Protocols for genotyping followed those described in Cacossi et al. (2019).

### Phylogenetic inferences and divergence times

To determine the origin and intraspecific divergence time among *Vriesea oligantha* populations, we conducted time calibration analyses in two steps. First, we performed a dating analysis of the Tillandsioideae subfamily, including six individuals of *V. oligantha* from different populations in a matrix with 171 species of Tillandsioideae and seven species from other subfamilies of Bromeliaceae using three plastidial (*matK, rpoB-trnC* and *ycf1*) and one nuclear (*phyC*) markers downloaded from Genbank (see Appendix S1 in Supporting Information). The phylogenetic tree for Tillandsioideae was inferred using the GTR+I+G site model, estimated in jModelTest2 (Darriba, Taboada, Doallo & Posada, 2012), assuming a Yule speciation prior, a lognormal prior distribution for calibration points and a normally distributed prior root set to mean=19 Myr, SD=1.25 under an uncorrelated lognormal relaxed clock model. The calibration age was extracted from Givnish et al. (2011).

Secondly, using the crown age obtained for the *Vriesea oligantha* clade, we ran a calibration approach to infer the dates of the intraspecific diversification of *V. oligantha*. We used samples from the 95 individuals described above, using seven other Tillandsioideae as outgroups. We used a normal distribution prior for the calibration point at the root, the GTR+I+G substitution rate model selected using jModelTest2 and a coalescent constant size random tree under an uncorrelated exponential relaxed-clock model.

Both steps were run in BEAST 1.10.4 and executed in CIPRES Portal (Drummond & Rambaut, 2007; Miller, Pfeiffer & Schwartz, 2010). Each analysis had four independent MCMC runs of 200 million generations sampled every 1,000. We used Tracer 1.10.4 to assess the convergence and effective sample size (ESS>200) for each run. As a burn-in, we excluded 10% of trees for each step, combining the remaining trees in LogCombiner. Maximum clade credibility trees for Tillandsioideae and *Vriesea oligantha* were inferred using TreeAnnotator and visualized in FigTree 1.4.4 and in the Interactive Tree of Life platform (https://itol.embl.de/).

### Genetic diversity

For cpDNA, we calculated the nucleotide diversity (π), haplotype diversity (Hd) and polymorphic sites (ss) per population in ARLEQUIN 3.5 (Excoffier & Lischer, 2010). A haplotype matrix was constructed based on ss detected, which were coded as single characters. To explore historical relationships among haplotypes, we built a median-joining-network (Bandelt, Forster & Rohl, 1999) on Network 5.0.1.1 (http://www.fluxus-engineering.com).

For nrSSR, putative clones were examined and removed using GenClone 2.0 (Arnaud-Haond & Belkhir, 2006). The number of alleles per locus (A), allelic richness per locus (AR), the observed (H_O_) and the expected (H_E_) heterozygosity, and the inbreeding coefficient (F_IS_) were estimated per population using GenAlex 6.5 (Peakall & Smouse, 2012). We evaluated the deviations from the Hardy-Weinberg equilibrium (HWE) per population and per loci using Genepop software v3.5 (Raymond & Rousset, 1995). Linkage disequilibrium between all pairs of loci was tested in FSTAT 2.9.3.2 (Goudet, 1995).

### Population structure and migration

For the population genetic structure of cpDNA, we employed a clustering analysis on GENELAND 4.0.7 (Guillot, Mortier & Estoup, 2005), testing 15 putative clusters. For nrSSR, we used STRUCTURE v.2.3.3 (Pritchard, Stephens & Donnelly, 2000) to assign individuals to genetic clusters (K) under the admixture model assuming independent allele frequencies. The number of K was set from 1 to 13, with 1,000,000 simulations for each K-value and a burn-in rate of 20%. The most probable number of K was examined using Structure Harvester v.6.0 (Earl & von Holdt, 2012), following the instructions of Evanno, Regnaut, & Goudet (2005).

Genetic differentiation was estimated using F-statistics (Weir & Cookham, 1984), based on cpDNA and nrSSR. Pairwise F_ST_ between localities were calculated using ARLEQUIN 3.5. We also conducted a molecular variance analysis (AMOVA) to evaluate the partition genetic variance in hierarchical models grouping by lineages obtained from the phylogenetic cpDNA tree by running 10,000 permutations between groups. AMOVA analyses were implemented on ARLEQUIN 3.5.

Recent migration events were estimated in BAYESASS 3.04 (Wilson & Rannala, 2003). Samples were run for 1.0×10^8^ interactions with a 10% burn-in, sampling every 1,000 interactions. Ancient migration events were estimated using MCMC and coalescence theory approach, implemented in MIGRATE 4.4.3 (Beerli & Felsenstein, 2001), testing two models, using the four phylogenetic lineages as groups: a panmictic model and a neighbor-only model, where migration was only allowed between adjacent groups. For both models, we used a Brownian motion model of mutation under the maximum-likelihood framework and 10 replicates of 1,000,000 runs with a 10% burn-in.

### Paleodistribution and ancestral population connections

To predict the current and paleodistributions (i.e., Mid-Holocene, MH, 6 kyr; Last Glacial Maximum, LGM, 21 kyr; and Last Interglacial Maximum, LIG, 120-140 kyr) of *Vriesea oligantha*, we conduct species distribution modeling (SDMs) based on 55 unique occurrence records using an ensemble approach that combined the results from six distinct modeling algorithms (see Appendix S2 for detailed analyses). From the estimated SDMs, we generated population connectivity maps by summing the least-cost path (LCP) among all populations (Chan, Brown & Yoder, 2011) to investigate ancestral suitable connections. We created a Friction Layer by inverting the SDM to a dispersal cost layer on ArcMap 10.5 (ESRI, 2016). Next, we calculated corridors that minimize the cost of dispersal between populations by following paths of lowest frictions. To do so, we used the “Least-Cost Corridors and Paths>Pairwise: All Sites” tool from SDMtoolbox 1.1a (Brown, 2014), implemented on ArcMap 10.5.

### Demographic reconstruction

To evaluate the demographic history of *Vriesea oligantha*, we used the cpDNA data to calculate the Fu’s Fs, Taijma’s D and Rozas’ R2 statistics and tested their departures from neutrality based on 100,000 coalescent simulations with DnaSP (Librado & Rozas, 2009). We also inferred changes in the species’ effective population size through time using a Coalescent Bayesian Skyline Plot (BSP) approach implemented in BEAST 1.10.4 and performed two independent MCMC runs of 200 million generations for each analysis, sampling every 2000 generations in the CIPRES. The BSP analysis was explored using the total cpDNA data set for each clade of the species tree under a strict molecular clock using the substitution rate interval inferred by the divergence time estimated for the Tillandsioideae subfamily (see Results below). Best substitution models for both markers were defined according to the Akaike Information Criterion (AIC) on jModelTest 2.0.

For nrSSR data, we estimated contemporary effective population sizes using NEestimator 2.1, with the Molecular Coancestry Method (Do et al., 2014). Putative recent bottleneck events were estimated using the “Wilcoxon sign-rank test’ with ‘Two phased mutation model’, with a total of 5,000 simulation iterations using BOTTLENECK 1.2.02 (Piry, Luikart & Cornuet, 1999).

### Roles of climate and geography on population structure

We first tested whether populations are differentiated following a model of isolation by distance by conducting a Mantel test (Wright, 1965) in R-package ‘adegenet’ with 10,000 permutations between geographic and genetic pairwise distances matrices, measured as F_ST_/(1-F_ST_). Then, we implemented a Bayesian generalized linear mixed modeling (GLMM) approach to test whether population genetic structure of *V. oligantha* is driven by isolation by environment (i.e., temperature and precipitation) and/or isolation by distance models (see Appendix S3 for detailed method).

## RESULTS

### Phylogenetic inferences and divergence times

The topology of the time-calibrated tree of Tillandsioideae indicates that the *Vriesea oligantha* complex is monophyletic (posterior probability, pp = 1.00) and belongs to Vrieseinae subtribe (Figure 2, S4.1, Table S5.1). This Bayesian inference indicated a nucleotide substitution rate mean of 0.0011135 Ma^−1^ (95% HPD=0.0009402-0.0012894). The estimated time to the most recent common ancestor (TMRCA) of *V. oligantha* was dated at 3.26 Myr (95% HPD=4.48-1.83 Myr) (Figure 2, Table S5.1). This tree shows four well-supported lineages in distinct geographic regions of the Espinhaço Range (Figure 2, S4.2). The Northern clade includes the JAC and MKA populations (pp=0.86, 0.702 Myr, 95% HPD=1.47-0.076 Myr) while Chapada Diamantina clade includes the MCH, MUC, RCO and ABA populations (pp=1.00) and it is estimated to be the oldest lineage (1.10 Myr, 95% HPD=1.17-1.04 Myr). Diamantina Plateau clade, includes the GMO, DIB, DIC, DIM, SGO and LIC populations (pp=0.98, 0.861 Myr, 95% HPD=1.06-0.494 Myr) and the Southern region of Espinhaço comprehends a clade with CIP and OUR populations (pp=0.99, 0.708 Myr, 95% HPD=0.999-0.274 Myr).

**Figure 2.**
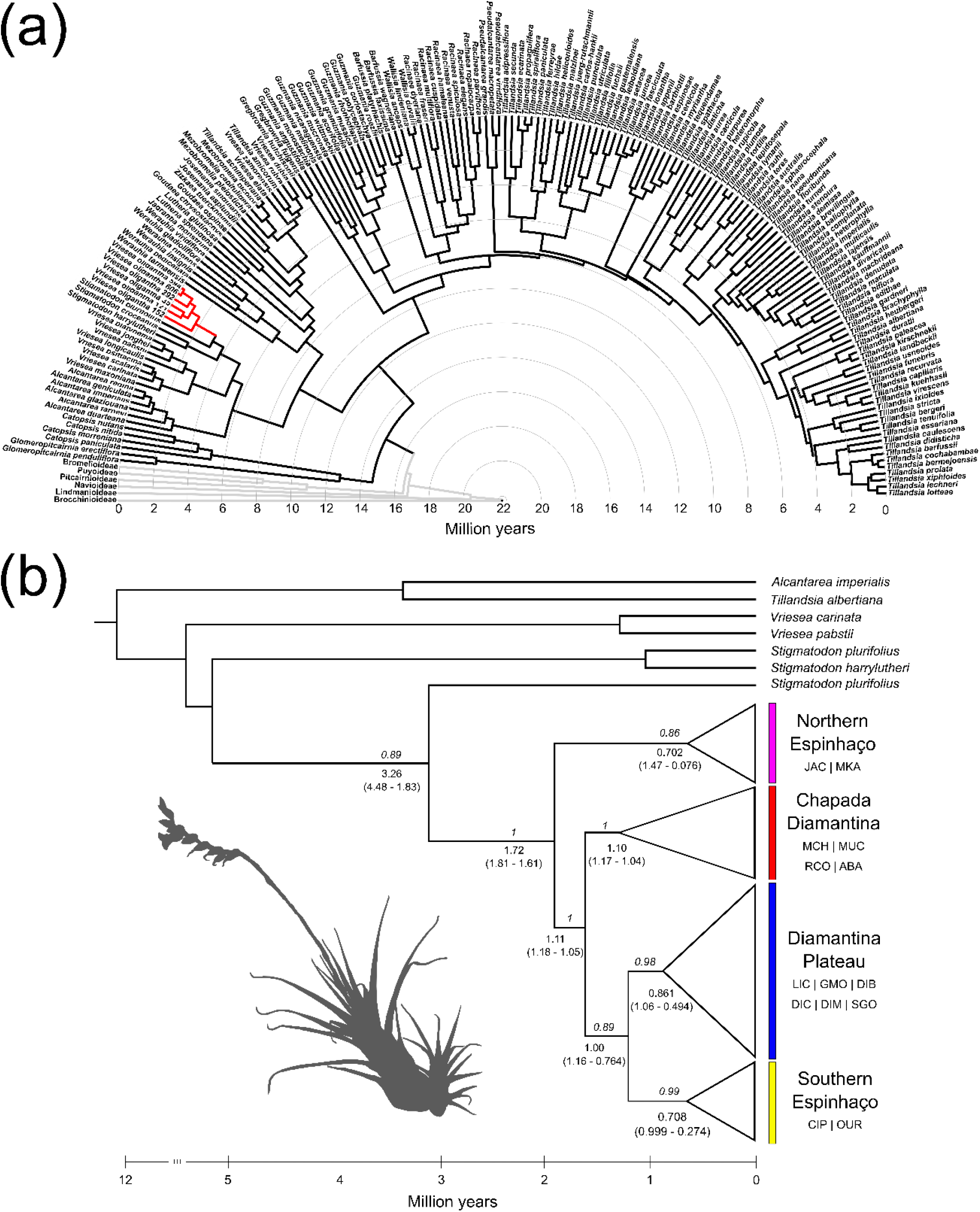
(a) Maximum clade credibility tree of Tillandsioideae. Branches in light grey are outgroups, branches in black are Tillandsioideae and branches in red are the *Vriesea oligantha* lineage. (b) Maximum clade credibility tree of *Vriesea oligantha*, depicting four lineages: Northern Espinhaço (pink), Chapada Diamantina (red), Diamantina Plateau (blue) and Southern Espinhaço. Bayesian posterior probabilities are in italics above each node; clade ages (in million years) are written under each node; values in parentheses are the 95% highest posterior density (HPD). A silhouette of *Vriesea oligantha* is in the inset.

### Genetic diversity

The final alignment matrix with consensus cpDNA sequences of *Vriesea oligantha* had 2,065 bp (*trnLF-trnF,* 753 and *ycf1,* 1312), showing 17 polymorphic sites and 22 haplotypes (Figure 1). The haplotype diversity per population ranged from zero to 0.785, and the nucleotide diversity from zero to 0.000761, with an overall haplotype and nucleotide diversity of 0.392 and 0.000280, respectively (Table 2). Few haplotypes are shared among populations, but the most frequent haplotype (H7) was found in five populations of the Diamantina Plateau clade, which also showed the highest values of nucleotide and haplotype diversity (Figure 1, Table 2).

**Table 2.**
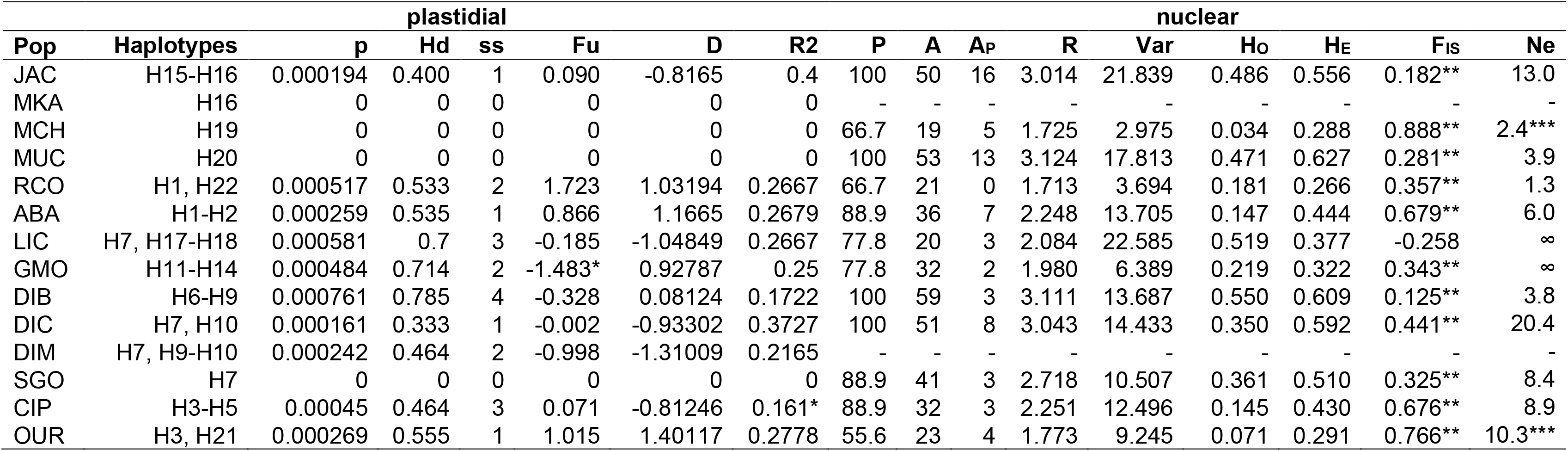
Genetic diversity and tests of neutrality of 14 populations of *Vriesea oligantha.*

We found a total of 162 alleles across loci and 67 were exclusive to single populations. Populations JAC and MUC presented the highest number of private alleles with 16 and 13 alleles, respectively (Table 2). Populations of *Vriesea oligantha* showed an averaged allelic richness ranging from 1.713 to 3.124, whereas variance in allele size ranged from 2.975 to 22.585 (Table 2). Observed and expected heterozygosity per population varied from 0.034 to 0.550 and 0.266 to 0.627, respectively. The mean inbreeding coefficient (F_IS_) was high and significant for most populations, with the exception of LIC population ranging from 0.125 in DIB and 0.888 in MCH (Table 2).

### Population structure

GENELAND results, based on cpDNA markers, indicated five clustering among populations of *Vriesea oligantha* (Figure 3a-e). Northern populations (JAC, MKA and MCH) form a single cluster, whilst populations from Chapada Diamantina (ABA, MUC and RCO) are clustered together. Diamantina Plateau populations (DIB, DIC, DIM, LIC and SGO) belong to a separate cluster while GMO constitute a single cluster. The Southern Espinhaço populations (CIP and OUR) form the fifth cluster. STRUCTURE clustering analysis (Figure 3f), based on nrSSR revealed a large number of genetic clusters among all populations (*k* = 11).

**Figure 3.**
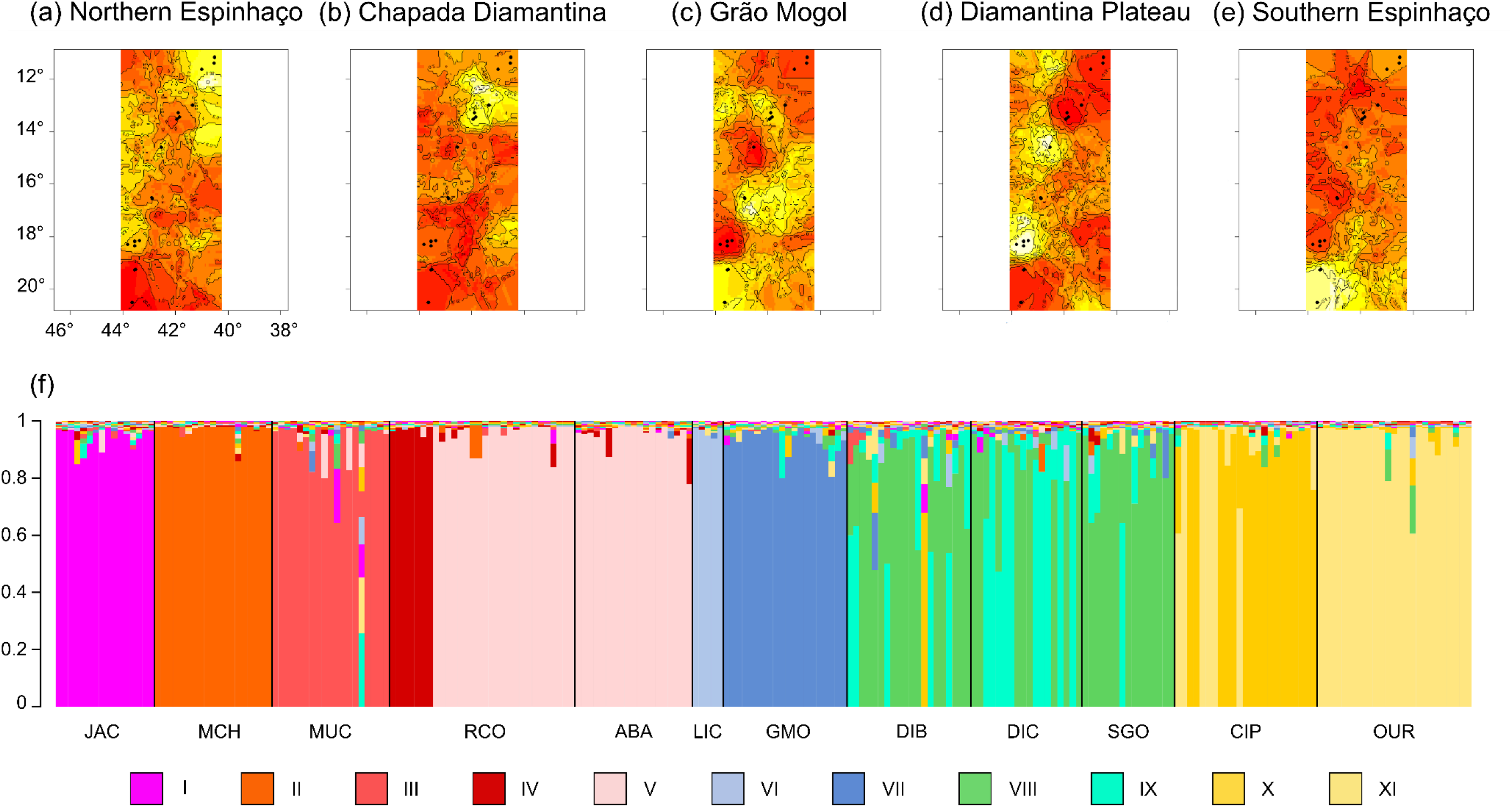
Clustering methods used for *Vriesea oligantha* populations based on cpDNA (a-e) and nrDNA (f). In GENELAND analysis (a-e), yellow areas are associated with a high probability of distinct populations belonging to each cluster (See text for population assignment). In STRUCTURE analysis (f), each color represents a given cluster (I-XI) and each bar corresponds to a single individual.

The pairwise F_ST_ values among populations were significant for almost all combinations, ranging 0.117-1.000 for cpDNA data and 0.007-0.794 for nrSSR data, indicating a widely heterogeneous structure between populations (Figure 4a-b, Tables S5.2-5.3). AMOVA showed high values of structure for both markers (cpDNA, 0.822; nrSSR, 0.432) and, hierarchically, cpDNA showed that the largest proportion of variance was confined within population (0.853, p<0.001) (Table 3).

**Figure 4.**
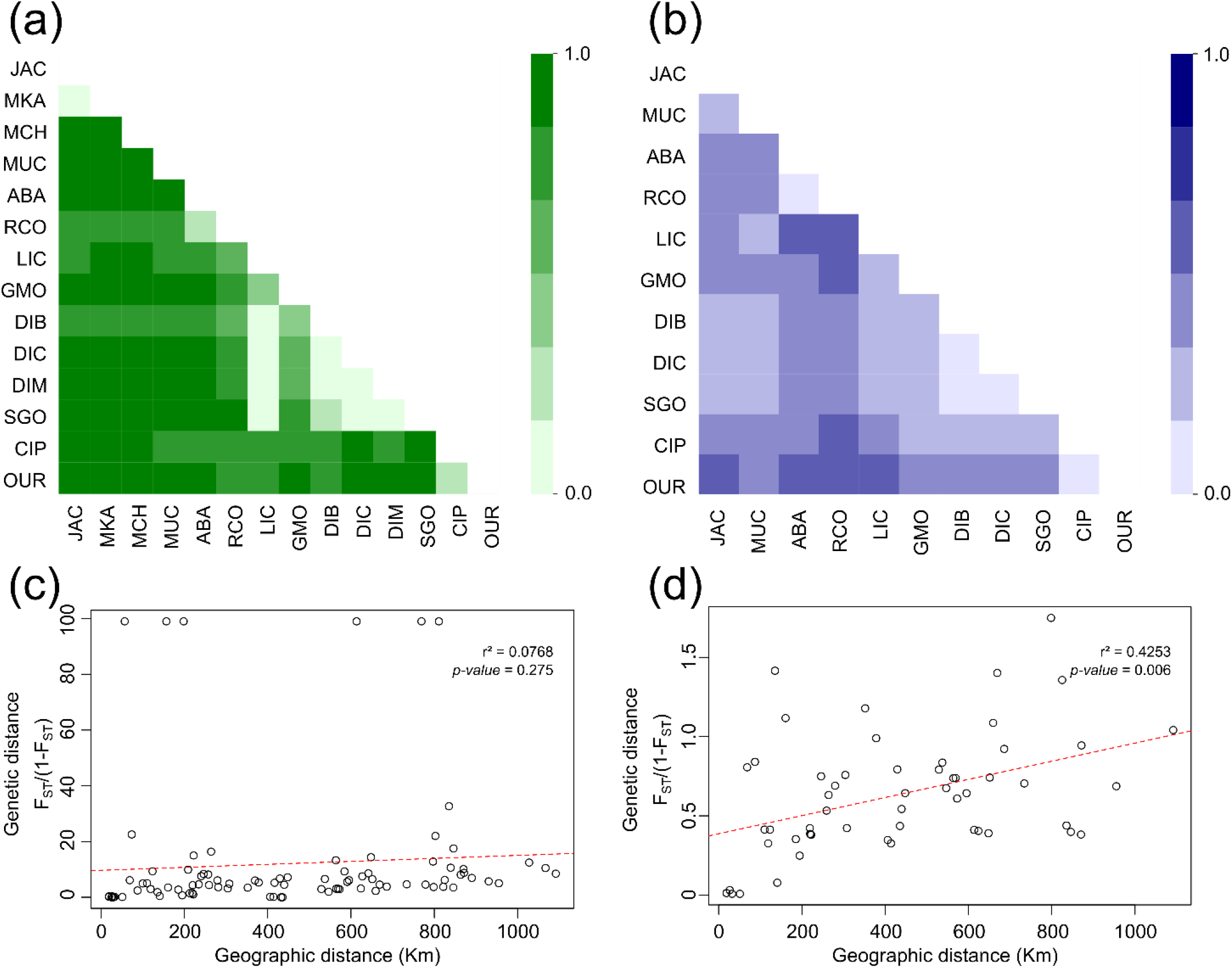
Pairwise F_ST_ heatmap plots and relationship between genetic and geographic distances, from plastidial (a, c) and nuclear (b, d) data.

**Table 3.** Analysis of molecular variance on cpDNA sequences and nrSSR, based on GENELAND and STRUCTURE, respectively. All values of *F*-statistics are significant (*p-value* < 0.001).

| Clustering method | Source of variation | df | Sum of Squares | Variance components | Variation (%) | *F*-statistics |
| --- | --- | --- | --- | --- | --- | --- |
| **cpDNA** |  |  |  |  |  |  |
| All populations | Among populations | 13 | 125.787 | 1.38841 | 82.2 | $F_{ST} = 0.822$ |
|  | Within populations | 81 | 24.297 | 0.29997 | 17.7 |  |
| Phylogenetic lineages | Among lineages | 3 | 95.702 | 1.35333 | 65.9 | $F_{CT} = 0.659$ |
| | Among populations within lineages | 10 | 30.085 | 0.39953 | 19.4 | $F_{SC} = 0.571$ |
| | Within populations | 81 | 24.297 | 0.29997 | 14.6 | $F_{ST} = 0.853$ |
|  | *Total* | 94 | 150.084 | 2.05283 | 100 |  |
| **nrSSR** |  |  |  |  |  |  |
| All populations | Among populations | 11 | 302.447 | 0.70207 | 43.2 | $F_{ST} = 0.432$ |
|  | Within populations | 446 | 410.732 | 0.92092 | 56.7 |  |
|  | *Total* | 457 | 88079.555 | 238.4214 | 100 |  |

Gene flow estimation from BAYESASS shows very few contemporary migration events, only among nearby populations (Table S5.4). Similarly, MIGRATE estimated few ancient events between the tested models (Table S5.5), where the migration only between adjacent groups model was better-supported than the panmictic model (Table S5.6).

### Palaeoclimatic distribution and ancestral population connections

Our species distribution models (SDMs) indicated that 120-140 kyr ago (LIG), high suitable areas for *Vriesea oligantha* were concentrated in sparse fragments, southwards from the current distribution, although mid-suitability would be observed all over the Espinhaço Range. This suggests that the species might have survived in small areas in northwards and westwards of its distribution during this period (Figure 5d). Later, as the climate got colder, during the LGM (21 kyr), suitable areas of *V. oligantha* might have increased, becoming progressively connected (Figure 5c). According to our SDMs, the *V. oligantha*’s distribution became more fragmented once again during the Mid-Holocene (6 kyr), with suitable areas in the Northern Espinhaço (Figure 5b). The current distribution mainly reflects areas of high altitude in the Espinhaço Range (Figure 5a).

**Figure 5.**
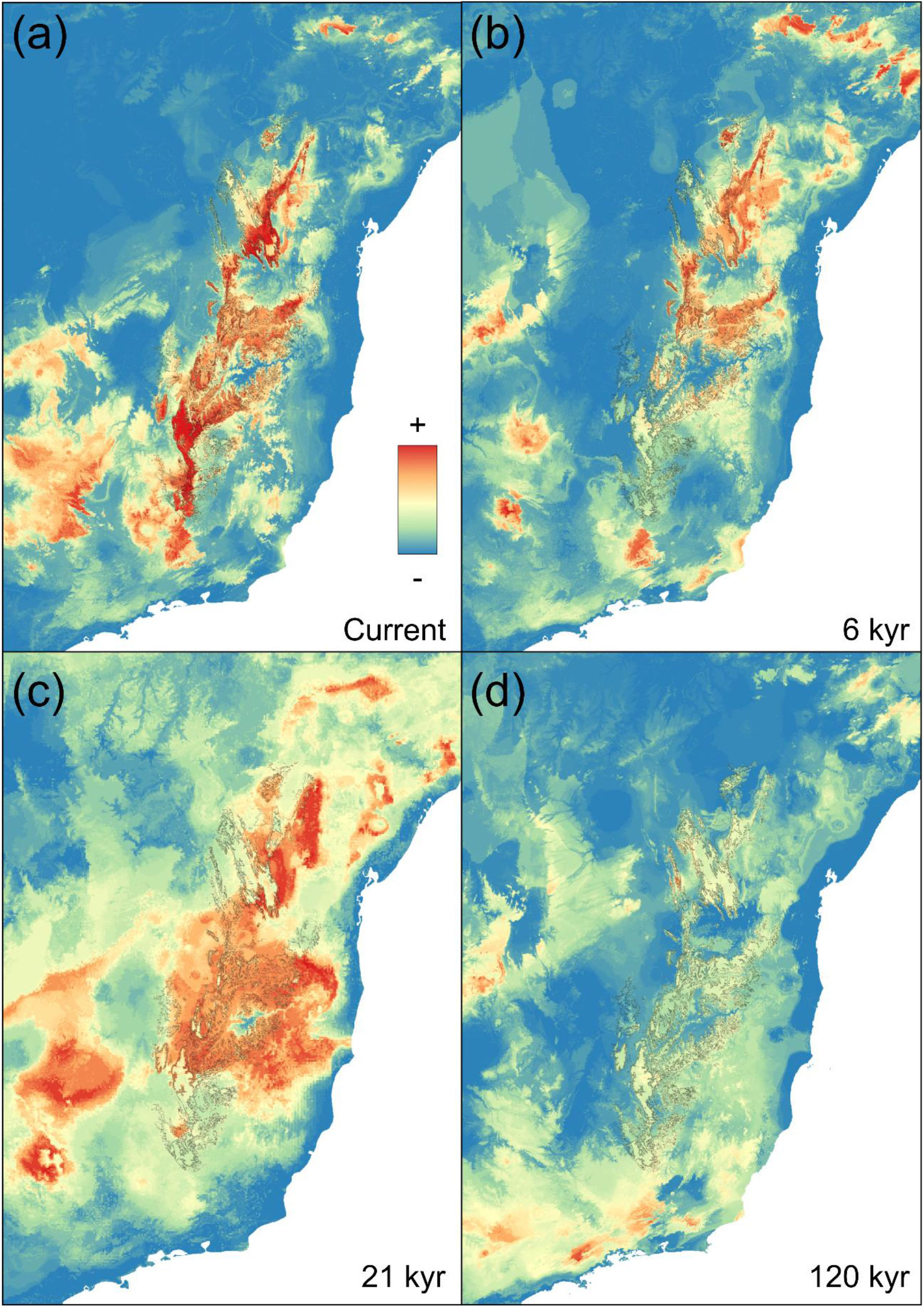
Ensemble distribution models for *Vriesea oligantha* under (a) Current, (b) Mid-Holocene (6 kyr), (d) Last Glacial Maximum (21 kyr) and (D) Last Interglacial Maximum (120 kyr) climatic conditions. The thin line represents the current delimitation of the Espinhaço Range.

Analysis of the least-cost path (LCP) across isolated populations revealed suitable areas of ancestral connection over all investigated periods, especially between the Chapada Diamantina and the Diamantina Plateau populations (Figure 6). However, those connections seem broader in the LIG (Figure 6d), while they become progressively narrow until the current climate conditions, with high suitability among populations in the Diamantina Plateau (Figure 6a-c). Populations from the northern and southern peripheries of the distribution presented lower connections with the central groups in all periods.

**Figure 6.**
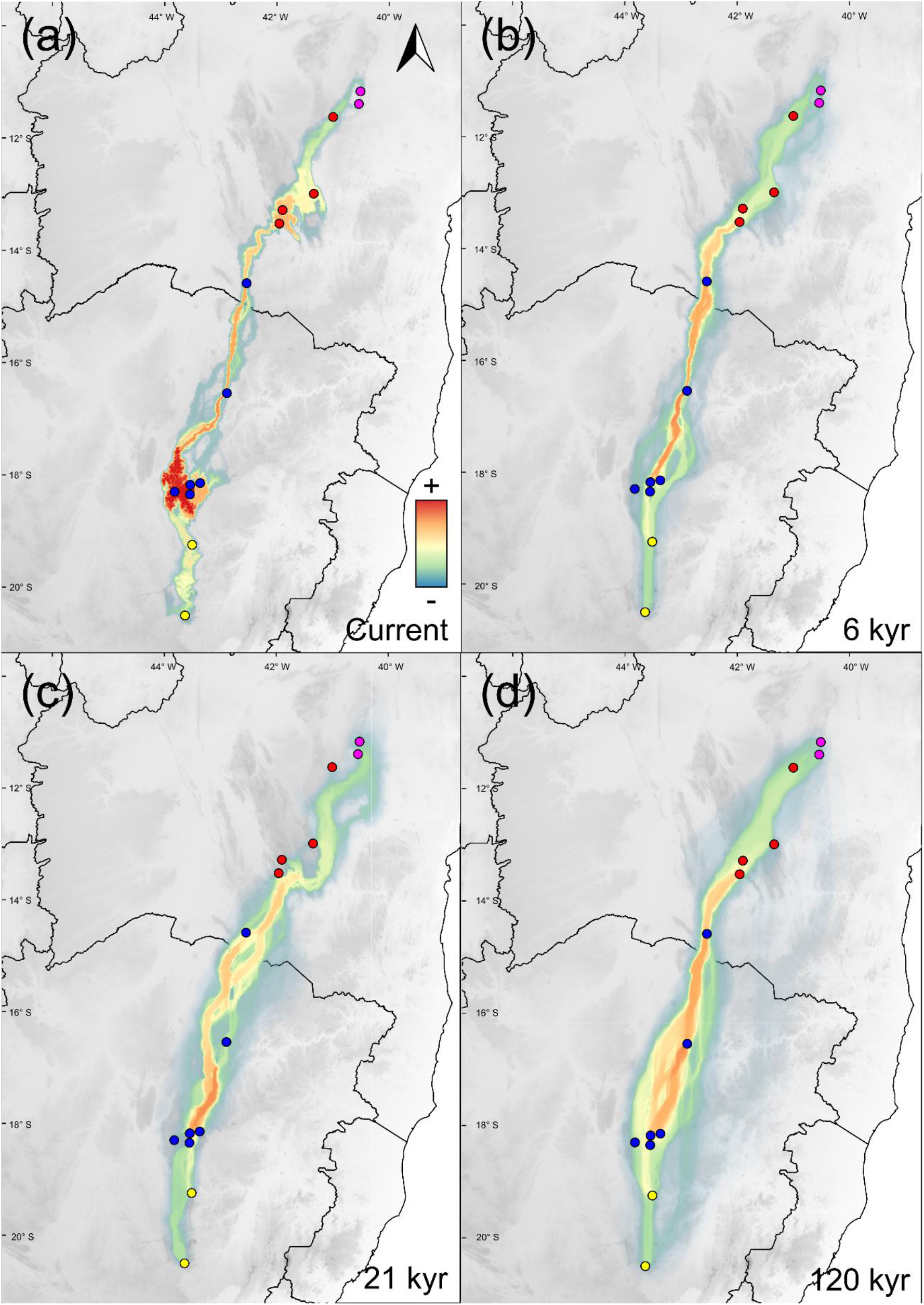
Potential dispersal corridors between *Vriesea oligantha* populations (circles, each color represents a lineage from phylogenetic tree) based on friction layers, across four period times. (a). Current; (b). Mid-Holocene (6 kyr); (d). Last Glacial Maximum (21 kyr); and (d). Last Interglacial Maximum (120 kyr). Altitude is represented by the grey background.

### Demographic analyses

All populations exhibited no departure from neutrality on demographic analyses, except for GMO (in Fu’s FS) and CIP (in Roza’s R2) (Table 2). Similarly, the Bayesian phylogenetic inference indicated a stable effective population size for the total dataset and to the four lineages during the last 100 kyr (Figure S4.3).

Contemporary effective population size ranged from 1.3 to 20.4, in populations RCO and DIC, respectively, while populations LIC and GMO showed infinite values (Table 2). Based on BOTTLENECK analysis, only MCH and OUR exhibited excess heterozygosity concerning the expectation under the TPM model (Wilcoxon test < 0.01, Table S5.7), evidencing that these two populations may have experienced population reduction.

### Relative roles of climate and geography

Mantel test showed a non-significant correlation between cpDNA differentiation and geographic distance (r^2^=0.07, *p*=0.275) (Figure 4c) but a significant correlation between nrSSR differentiation and geographic distance (r^2^=0.42, *p*=0.006) (Figure 4d), indicating isolation-by-distance in the nuclear genome (Diniz-Filho et al., 2013). Our GLMM analysis indicated the influence of both environmental and geographical distances on the genetic structure of *Vriesea oligantha*. Contrasted with other predictors, the model including both isolation by geographic distance (IBD) and by environment (IBE) received the highest DIC support for both plastidial and nuclear genomes (Table 4).

**Table 4.** Results of generalized linear mixed models (GLMM) testing the influence of geographic distance and climatic differences, based on 19 bioclimatic variables, on the genetic divergence among populations of *Vriesea oligantha*. The best model for each data set is highlighted in bold.

| Model | cpDNA |  |  | nrSSR |  |  |
| --- | --- | --- | --- | --- | --- | --- |
| | DIC | ΔDIC | DIC weight | DIC | ΔDIC | DIC weight |
| Null | 810.1204 | 9.6164 | 0.0077 | 113.2596 | 9.5066 | 0.0045 |
| Geography | 811.7689 | 11.2648 | 0.0034 | 106.2901 | 2.5372 | 0.1461 |
| Climate | 806.3522 | 5.8482 | 0.0504 | **103.7529** | **0.0000** | **0.5196** |
| Geography + Climate | **800.5041** | **0.0000** | **0.9386** | **104.6625** | **0.9096** | **0.3297** |

## DISCUSSION

### Diversification of *Vriesea oligantha*

The divergence age between *Vriesea oligantha* and its outgroup dated back to the Pliocene-Pleistocene transition (3.26 Myr), as previously inferred in phylogenies of *Vriesea* (Kessous et al., 2019; Machado et al., 2019). This period is marked by a decrease in global temperature and the beginning of glaciations in the Northern Hemisphere (Prell, 1984). In this context, *V. oligantha* is a remarkable model to test effects of climatic oscillations, since this species remained consistent over the whole Pleistocene.

Our results show that the intraspecific divergence events of *Vriesea oligantha* are older than those associated with the latest Pleistocene climatic oscillations and cannot be explained solely by the late-Quaternary refugia hypothesis (Haffler, 1969; Rull, 2011). Altogether, extreme amplitude in climatic fluctuations were remarkable events in the mid-Pleistocene (Lisiecki & Raymo, 2007), when *V. oligantha* lineages diversified, supporting the view that climate fluctuations from the early-Quaternary are key components for understanding population differentiation processes and, eventually, the role of the environment in speciation events of Neotropical lineages (Silva, Antonelli, Lendel, Moraes & Manfrin, 2018). In agreement, our demographic analyses were unable to demonstrate significant changes in the effective population size over the past thousands of years, a possible consequence of little or no relevance of the late-Quaternary events on population differentiation of this species. However, demographic analyses usually assume panmictic populations, leading to misinterpretations about demographic changes in highly structured populations, as in *V. oligantha* (Heller, Chikhi & Siegismund, 2013). Another caveat of demographic changes is the scarcity of polymorphisms in our genetic markers, a common and intrinsic issue found in Bromeliaceae (Versieux et al., 2012). In fact, diversification events throughout the Quaternary are also congruent with other extant lineages from distinct Neotropical montane formations, as in the Espinhaço itself (e.g., Barres et al., 2019; Bonatelli et al., 2014; Chaves et al, 2019; Nascimento et al., 2018; the páramos (e.g., Hughes & Atchison, 2015; Madriñán, Cortés & Richardson, 2013) and the pantepuis (e.g., Rull et al, 2020; Salerno et al., 2012), reinforcing the importance of climate changes throughout the whole Pleistocene in the diversification of the montane biota in the Neotropics.

### Phylogeography of *Vriesea oligantha*: insights from micro to macroevolution in the Espinhaço Range

The phylogeographic results of *Vriesea oligantha* showed a remarkable congruence between the structure of different populations and the bioregionalization proposed by Colli-Silva et al. (2019), based on patterns of plant endemism in the Espinhaço Range. Similar biogeographic patterns were also pointed out by several other studies using floristic and faunistic composition and endemicity indexes (e.g., Bitencourt & Rapini, 2013; Bünger, Stehmann & Oliveira-Filho, 2014; Campos, Freire Moro, Funk & Roque, 2019; Chaves et al., 2015) and phylogenetic histories (e.g., Chaves et al., 2019; Ribeiro et al., 2014). Taken together, these studies suggest that particular drivers are leading the evolutionary history, community assembling, and physiognomy patterns of distinct areas in the Espinhaço Range (Zappi, Moro, Meagher, & Nic Lughadha, 2017).

Our results support the congruency between the Northern Espinhaço and the Chapada Diamantina lineages and clustering analysis within the Chapada Diamantina province (Colli-Silva et al., 2019). Populations from the Diamantina Plateau clade mainly mirrors the Southern Espinhaço province, particularly the Diamantina Plateau district. However, the GMO population is a noteworthy exception. In fact, as evidenced by GENELAND and STRUCTURE, GMO remained distinct on a single cluster, unarguably fitting into the Grão-Mogol district. The particularity of this district has already been reported elsewhere (e.g., Echternacht et al. 2011; Pirani, Mello-Silva & Giulietti, 2003), where its discontinuity seems to be the factor that leads to the singularity of these mountains. Another relevant factor is linked to its location as an ecotone between the Cerrado and Caatinga phytophysiognomies, affecting the dynamics of the biological community of this region and resulting in a particular evolutionary history.

The fourth estimated clade, the Southern Espinhaço, includes CIP and OUR populations, the latter coinciding with the Iron Quadrangle district, the southernmost bioregion of the Espinhaço. However, we could not directly link this clade with the Iron Quadrangle district, since its diagnostic characteristic is the presence of ironstone outcrops, while CIP and OUR are associated with quartzitic soils (Saadi, 1995). Despite the lack of putative soil differences, the relative proximity of CIP and OUR with the Iron Quadrangle district may promote the divergence of the Southern Espinhaço clade due to biotic interactions and constraints fostered by the singular environment and community of the Iron Quadrangle (Jacobi, do Carmo, Vincent & Stehmann, 2007; Zappi et al., 2017). Further analyses exploring the role of ecological interactions and how they affect speciation among communities could improve this hypothesis (Johnson & Stinchcombe, 2007)

### Microevolutionary processes of *Vriesea oligantha* are driven by geographic distance and climate gradient

Our GLMM analysis demonstrates that populations of *Vriesea oligantha* are structured both by IBD and IBE. Indeed, both pollen and seed dispersal might be of great importance to explain the geographic component of the genetic structure pattern observed in nuclear and plastidial markers. However, assuming maternal inheritance of plastidial DNA and biparental inheritance of nuclear DNA as a rule for Angiosperm (Ennos, 1994), our genetic differentiation results indicate that gene flow via seeds is comparatively less efficient than gene exchange via pollen, in agreement with non-significant IBD obtained for the plastidial marker in the Mantel test. The discrepancy between the two markers may indicate the role of pollinators in maintaining the cohesion between close populations, since seeds are poorly dispersed, a common trend in other plant lineages of the Espinhaço Range (Silveira et al. 2020). Small and isolated populations may experience genetic depletion and local extinction, but if these populations persist, they might differentiate until the reproduction incompatibility and/or ecological divergence evolve, critical steps towards speciation (Harvey, Singhal & Rabosky, 2019). Thus, extant organisms with low dispersion that inhabit small and fragmented environments, such as the mountains of the Espinhaço Range, could be more prone to experience speciation events (Kisel & Timothy, 2010) and the described microevolutionary processes of *V. oligantha* might be the first steps of its diversification (Pinheiro et al., 2013).

Climatic variables along the Espinhaço Range was also a determinant component in the current genetic structure of the populations of *Vriesea oligantha.* The Northern Espinhaço climate is markedly drier and hotter, with long periods of low (or even absence of) precipitation while the climate of the mid-south Espinhaço has milder temperatures and higher humidity levels (Giulietti, Menezes, Pirani, Meguro & Wanderley, 1987; Zappi et al., 2003). Such variation has the potential to rapidly influence evolution by triggering ecological divergence (Campbell & Powers, 2015), especially in a scenario with reduced gene flow as observed in *V. oligantha*. Ancient climatic changes have a potential to strongly impact montane biological communities, as also reported in the Andes (Flantua et al., 2019). Paleopalinological studies of the Espinhaço have evidenced herbaceous vegetation expansion in the past towards lowlands (Barros, Lavarini, Lima & Magalhães-Júnior, 2011; Behling, 2002; Horák-Terra et al., 2015), supporting the putative role of ancestral climatic oscillations in the macroevolutionary patterns of endemic lineages in the Espinhaço Range (Vasconcelos et al., 2020). Nevertheless, considering the lower elevation of the Espinhaço Mountains (700-2,072 m) compared with the Andes (4,000-6,961 m), we expect the impacts of Pleistocene climate changes on *V. oligantha* diversification to be less associated with local extinctions due to glaciation events since there are few or no evidence that these mountaintops were once completely frozen (Luiz, Carneiro & Benitez, 2001). Thus, as estimated by the SDMs and the corridors between populations, events of secondary contact, hybridization and ecological interactions with other organisms are more likely to have promoted the current phylogeographic pattern in *V. oligantha*.

The evolution of species-rich biotas is remarkably known for its complex drivers (Antonelli et al., 2018; Rull, 2011) and other triggers might also help understanding the intra-specific lineage diversification of organisms from montane systems worldwide, such as edaphic adaptations (Alcantara et al., 2018), pollination strategies (Franceschinelli, Jacobi, Drummond & Resende, 2006) and niche conservatism (de Mattos, Morellato, Camargo & Batalha, 2019). In the future, the relative role of such processes could be tested by incorporating association analyses of the entire or partial genomes from multiple species with environmental datasets and functional ecological traits under a comparative framework.

### Microevolutionary processes as a proxy for understanding macroevolutionary patterns

We showed that microevolutionary processes underlying the phylogeographic patterns of *Vriesea oligantha* are a consequence of IBD and IBE throughout its distribution. Considering the assumption that population differentiation is the basic mechanism of speciation, the concordant patterns between the diversification of *V. oligantha* and the Espinhaço biogeography generate powerful insights into how climatic variables and limited gene flow shaped early stages of macroevolutionary patterns. Studies rarely attempt to fill the gap between micro and macroevolutionary approaches and studies like ours are necessary to pave the way on the initial effects of population differentiation that ultimately lead to the origin of new species and spatial patterns of biodiversity distribution. Additional evidence using distinct organisms could also provide contrasting examples of how microevolutionary processes act and translate into the current biogeographic patterns of tropical montane biotas.

## Acknowledgments

We thank Dr. Kléber Resende (CENA, USP) for fieldwork assistance.

Funding by Coordenação de Aperfeiçoamento de Pessoal de Nível Superior (CAPES) (Grant numbers: 88881.128215/2016-01 and PDSE 88881.190071/2018-01, PROAP-2015/CAPES/PPGCB-BV), FAPESP (2018/07596-0), CNPq (produtividade 300819/2016-1 and 305398/2019-9), CNPq (produtividade 304778/2013-3, 303794/2019-4 and Universal Project 455510/2014-8).

## DATA AVAILABILITY STATEMENT

We will submit the DNA sequences used in this work on GenBank (NCBI) after the publication acceptance.

## BIOSKETCH

**Marcos Vinicius Dantas-Queiroz** is a PhD candidate at São Paulo State University, Brazil. His broader research interests include biogeography, phylogeography, ecology and evolution of Neotropical plants, with emphasis on understanding the processes responsible for origin and diversification of species. All authors share a common interest in understanding the processes and mechanisms responsible for the generation and maintenance of biodiversity using mainly bromeliads as Neotropical models to reach this goal.

## AUTHOR CONTRIBUTIONS

M.V.D.Q., C.P.S. and L.M.V. conceived the ideas and designed the study; M.V.D.Q., T.C.S. and L.M.V. conducted fieldwork and collected the data. M.V.D.Q., T.C.S., B.S.S.L., C.J.N.C., T.N.C.V. and C.P.S. analyzed the data. M.V.D.Q. wrote the manuscript and all authors helped improve the final version. The authors declare no conflict of interest.

