## Appendix S1-S5 for "Underlying microevolutionary processes parallel macroevolutionary patterns in ancient Neotropical Mountains": Appendix S2-S3.docx

**Appendix S2 -** **Material and Methods of Paleodistribution and ancestral population connections**

To predict the current and paleodistributions of *V. oligantha*, we conduct species distribution models (SDMs). We first listed 55 unique occurrence records of *V. oligantha* from field trips and SpeciesLink database (http://splink.cria.org.br), which we mapped in a squared geographical grid of c.a. 3,500,00 km² and 30’’ of resolution (ca. 1 km²). To characterize the background environmental conditions, we selected the following five less correlated bioclimatic variables of the 19 available in Worldclim database (http://worldclim.org; (Hijmans, Cameron, Parra, Jones & Jarvis, 2005), through a factorial analysis, using ‘psych’ R package (Revelle, 2015): annual mean temperature (°C), temperature seasonality (°C), temperature annual range (°C), annual precipitation (mm), and precipitation of driest month (mm). To predict the current potential distribution of *V. oligantha*, we adopted six distinct algorithms: (i) Bioclim (Nix, 1986), (ii) Mahalanobis distance (Farber & Kadmon, 2003), (iii) Domain (Gower distance; Carpenter, Gillison & Winter (1993), (iv) Maximum Entropy (MAXENT; Phillips & Dudík (2008), (v) Support Vector Machines (SVM; Tax & Duin (2004), and (vi) Random Forest (Breiman, 2001).

To evaluate the predictions of potential *V. oligantha* distribution, we randomized the occurrence records in two subsets (75 and 25 % of train and test records, respectively) and estimated the true skill statistics (TSS; Allouche, Tsoar & Kadmon (2006). From a range of -1 (worse) to 1 (ideal), the acceptable models showed a TSS > 0.5 (Allouche et al., 2006). To predict the effects of past climatic oscillations on *V. oligantha* distribution, we projected the models onto the paleoclimatic scenarios simulated by the Community Climate System Model (CCSM4; Hijmans et al. 2005) for the Mid-Holocene (MH, 6 kya), Last Glacial Maximum (LGM, 21 kya), and Last Interglacial (LIG,120-140 kya; Otto-Bliesner, Marshall, Overpeck, Miller & Hu, 2008). To increase the reliability of potential distribution predictions, we employed an ensemble approach of all models using the average suitability weighted by the TSS value of each model (see Barry & Elith (2006); Diniz-Filho et al. (2009).

**Appendix S3 - Material and Methods of Roles of climate and geography on population structure**

We implemented a Bayesian generalized linear mixed modeling (GLMM) approach to test whether genetic structure among *V. oligantha* populations is driven by environmental variables, i.e. temperature and precipitation, and/or geographic distance among populations. We used the pairwise linear F_ST_ matrices [measured as F_ST_/1(1-F_ST_)] from both cpDNA and nrSSR as the response variable. As predictor variables, we tested a matrix of the natural logarithm of geographic distances among populations to assess the effect of isolation by distance (IBD) and a matrix of Euclidean distances along the first and second axis of a principal component analysis (PCA) of 19 WorldClim variables (Hijmans et al. 2005) at a resolution of 30 arc seconds, to access the effect of isolation by environment (IBE). The first and second axes of the PCA accounted for 62.77 and 21.64% of the data variance. This first axis was mostly correlated with Precipitation of Wettest Quarter, Precipitation of Wettest Month and Minimal Temperature of Coldest Month, while the second axis were more associated with the Precipitation of Driest Month and Precipitation of Driest Quarter.

We confronted models including only IBD, only IBE or both IBD and IBE as predictors against a null model. We performed the analysis with the R packages MCMCGLMM (Hadfield, 2010), following scripts available in Lexer et al. (2014). We ran 2,000,000 MCMC iterations with a 500,000 burn-in with a thinning interval of 750, under standard priors. The lack of independence between pairs of populations was accounted by fitting a multiple membership model. Chain convergences were checked using ‘CODA’ R packages. We used the Deviance Information Criterion (DIC) to compare models and determine the role of IBD versus IBE in the population structure of populations of *V oligantha*.
