## Appendix S1-S5 for "Underlying microevolutionary processes parallel macroevolutionary patterns in ancient Neotropical Mountains": Appendix S4.docx

**Appendix S4 – Supplementary Figures**

Figure S4.1 - Phylogenetic tree of Tillandsioideae. Branch colors are the posterior probability of each node, varying from 0 (red) to 1 (blue). The bottom scale is in million years.

Figure S4.2 – Phylogenetic tree of *Vriesea oligantha*. Branch colors are the posterior probability of each node, varying from 0 (red) to 1 (blue). The bottom scale is in million years

Figure S4.3 – Bayesian Skyline Plots (ESP) from each of the phylogenetic lineages (A-D) and for the total dataset of *Vriesea oligantha*, with 95% HPD intervals (thinner lines). The y-axis indicates population size; the x-axis indicates the time in millions of years

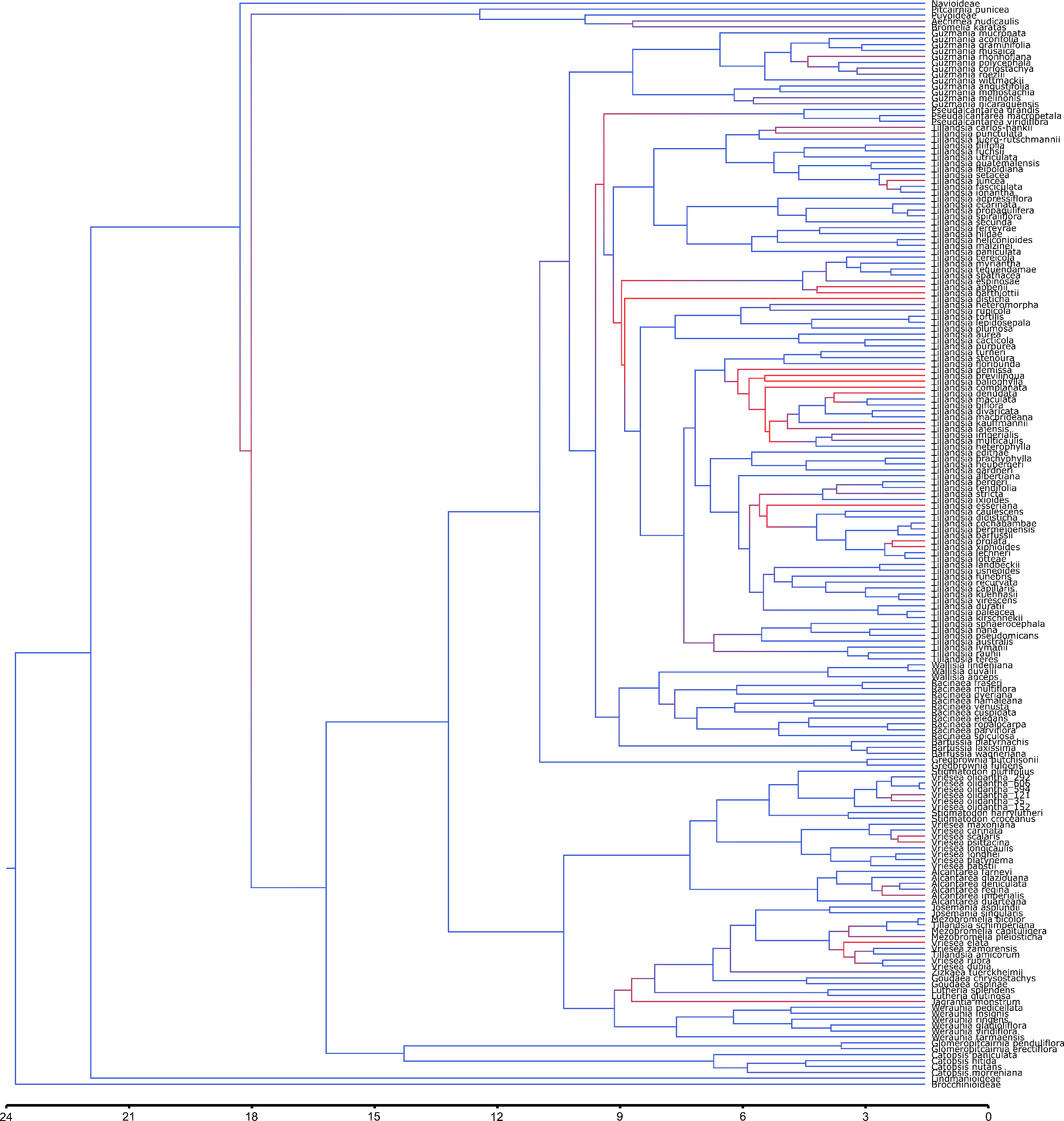

Figure S4.1 - Phylogenetic tree of Tillandsioideae. Branch colors are the posterior probability of each node, varying from 0 (red) to 1 (blue). The bottom scale is in million years.

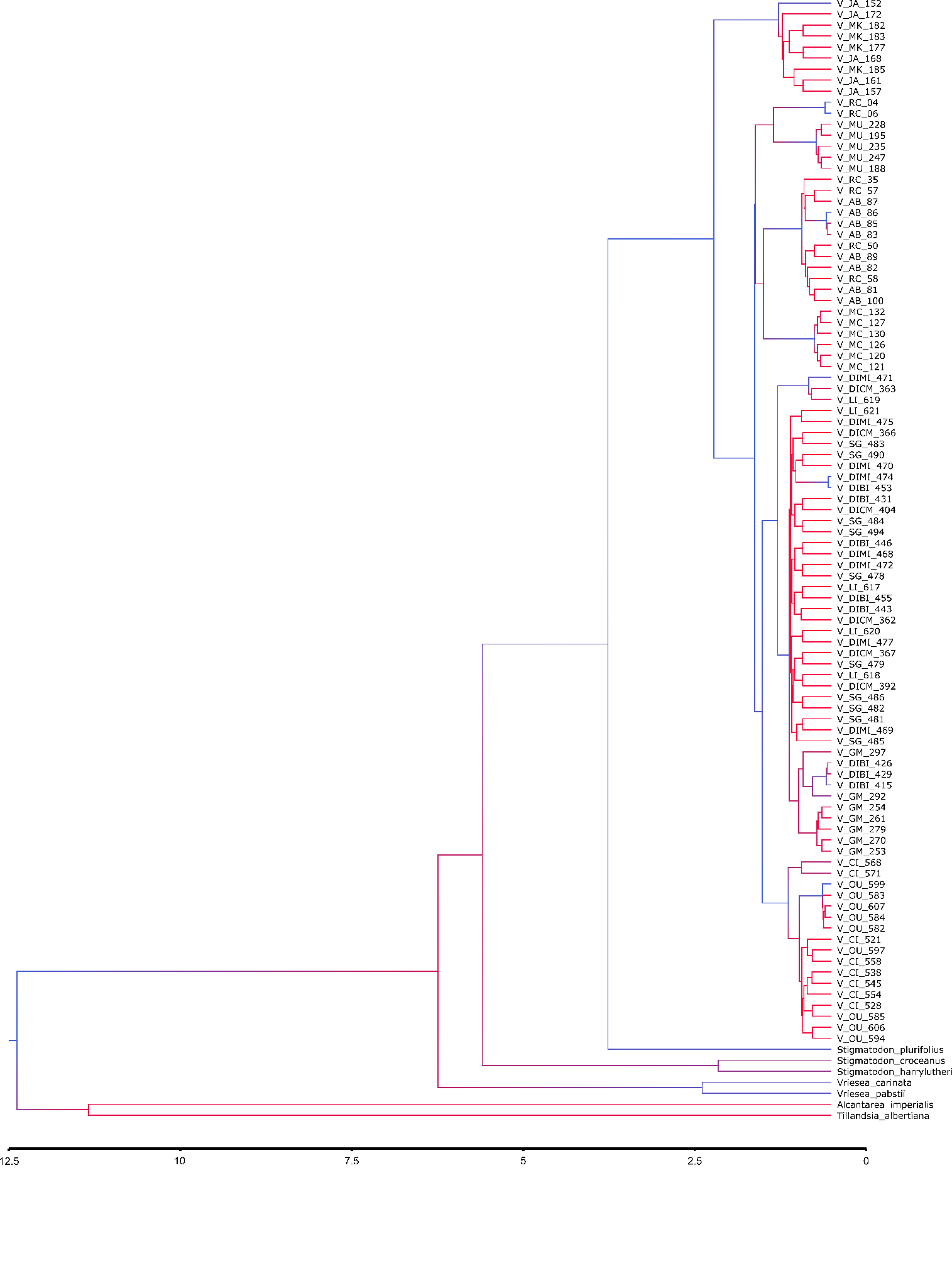

Figure S4.2 – Phylogenetic tree of *Vriesea oligantha*. Branch colors are the posterior probability of each node, varying from 0 (red) to 1 (blue). The bottom scale is in million years

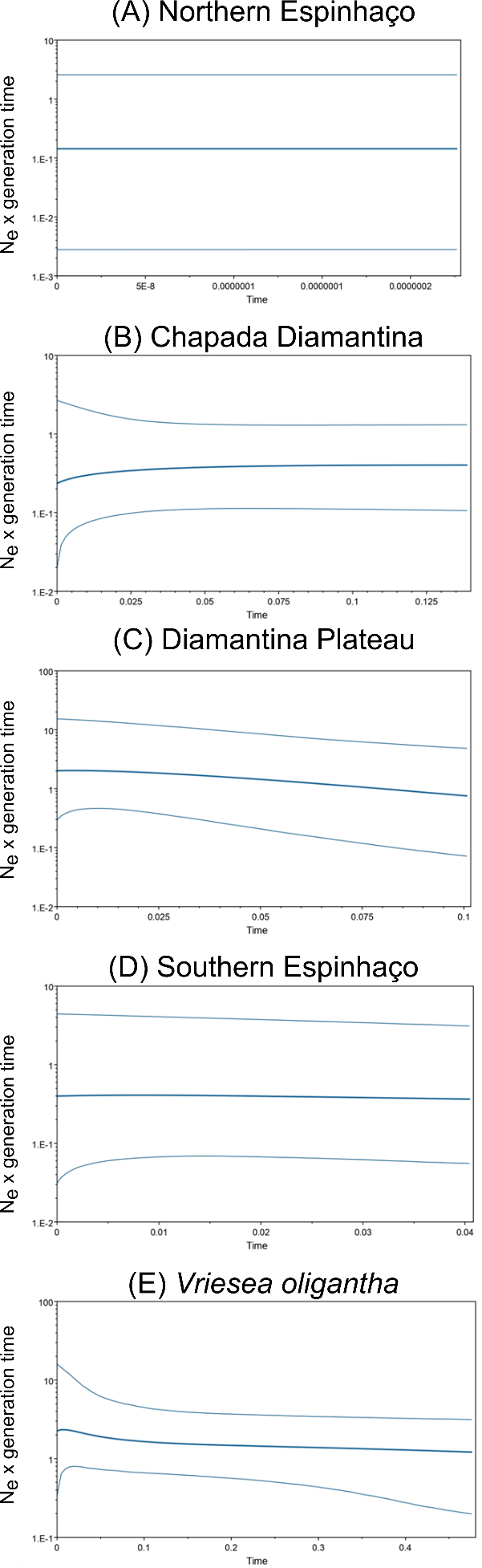

Figure S4.3 – Bayesian Skyline Plots (ESP) from each of the phylogenetic lineages (A-D) and for the total dataset of *Vriesea oligantha*, with 95% HPD intervals (thinner lines). The y-axis indicates population size; the x-axis indicates the time in millions of years
