## Appendix S1-S5 for "Underlying microevolutionary processes parallel macroevolutionary patterns in ancient Neotropical Mountains": Appendix S5.docx

**Appendix S5 – Supplementary Tables**

Table S5.1 - Estimated ages (Mya) in Tillandsioideae using a relaxed-clock model approach. The values in brackets are the 95% highest posterior density (HPD). Tribes and subtribes following Barfuss et al. 2016. For population names, see Table 1.

Table S5.2 - Pairwise F_ST_ among 14 populations of Vriesea oligantha based on cpDNA sequences. All significant values are in bold.

Table S5.3 - Pairwise F_ST_ among 12 populations of Vriesea oligantha based on nrDNA microsatellites. All significant values are in bold.

Table S5.4 - Contemporary migration rates estimated in BAYESASS among 12 populations of *Vriesea oligantha*. Estimated migration rates > 0.10 are displayed in red.

Table S5.5 -Posterior distribution values for the *Neighbor* groups only, estimated on Migrate.

Table S5.6 – Probability of the data of the two tested models (marginal likelihood) estimated on MIGRATE.

Table S5.7 – Probability of population reduction, estimated *with* BOTTLENECK 1.2.02, using all loci fitting the two phases model (TPM).

Table S5.1 - Estimated ages (Mya) in Tillandsioideae using a relaxed-clock model approach. The values in brackets are the 95% highest posterior density (HPD). Tribes and subtribes following Barfuss et al. 2016. For population names, see Table 1.

| **Taxon** | **Crown age (Mya)** | **bpp** |
| --- | --- | --- |
| Tillandsioideae | 14.6 (16.0 – 13.2) | 1 |
| Glomeropitcairnieae + Catopsideae | 12.7 (14.8 – 10.4) | 0.99 |
| Vrieseeae | 8.8 (10.3 – 7.3) | 1 |
| Cipuropsidinae | 7.5 (9.1 – 6.1) | 1 |
| Vrieseinae | 5.7 (10.3 – 7.3) | 1 |
| *Alcantarea* | 2.6 (3.8 – 1.4) | 1 |
| *Vriesea* | 3.0 (4.2 – 1.9) | 1 |
| *Stigmatodon + Vriesea oligantha* | 3.8 (5.2 – 2.5) | 0.99 |
| *Vriesea oligantha* | 1.7 (2.6 – 0.9) | 1 |
| Tillandsieae | 9.4 (10.7 – 8.0) | 1 |
| *Guzmania* | 7.1 (8.6 – 5.6) | 1 |
| *Gregbrownia* | 1.4 (2.6 – 0.4) | 1 |
| *Barfussia* | 1.7 (2.9 – 0.7) | 1 |
| *Wallisia* | 2.3 (4.0 – 0.9) | 1 |
| *Racinaea* | 6.1 (7.3 – 4.8) | 0.87 |
| *Pseudoalcantarea* | 2.9 (5.0 – 1.2) | 1 |
| *Tillandsia* | 7.6 (8.8 – 6.3) | 0.96 |
| **Clade** |  |  |
| Northern Espinhaço (JAC \| MKA) | 0.702 (1.473 – 0.076) | 0.86 |
| Chapada Diamantina (MCH \| MUC \| RCO \| ABA) | 1.108 (1.177 – 1.040) | 1 |
| Diamantina Plateau (LIC \| GMO \| DIB \| DIC \| DIM \| SGO) | 0.861 (1.065 – 0.494) | 0.98 |
| Southern Espinhaço (CIP \| OUR) | 0.708 (0.999 – 0.274) | 0.99 |

bpp = Bayesian posterior probability

Table S5.2 - Pairwise F_ST_ among 14 populations of Vriesea oligantha based on cpDNA sequences. All significant values are in bold.

|  | **JAC** | **MKA** | **MCH** | **MUC** | **ABA** | **RCO** | **LIC** | **GMO** | **DIB** | **DIC** | **DIM** | **SAI** | **CIP** | **OUR** |
| --- | --- | --- | --- | --- | --- | --- | --- | --- | --- | --- | --- | --- | --- | --- |
| **JAC** | **0** |  |  |  |  |  |  |  |  |  |  |  |  |  |
| **MKA** | -0.05263 | **0** |  |  |  |  |  |  |  |  |  |  |  |  |
| **MCH** | **0.95752** | **1** | **0** |  |  |  |  |  |  |  |  |  |  |  |
| **MUC** | **0.9375** | **1** | **1** | **0** |  |  |  |  |  |  |  |  |  |  |
| **ABA** | **0.85853** | **0.89116** | **0.90829** | **0.85948** | **0** |  |  |  |  |  |  |  |  |  |
| **RCO** | **0.7611** | **0.78378** | **0.82222** | **0.70944** | 0.25011 | **0** |  |  |  |  |  |  |  |  |
| **LIC** | **0.81818** | **0.83871** | **0.85799** | **0.8125** | **0.77866** | **0.6476** | **0** |  |  |  |  |  |  |  |
| **GMO** | **0.86663** | **0.8826** | **0.90267** | **0.87069** | **0.84106** | **0.77451** | **0.49947** | **0** |  |  |  |  |  |  |
| **DIB** | **0.77809** | **0.78872** | **0.8201** | **0.75904** | **0.74264** | **0.66231** | 0.12173 | **0.40906** | **0** |  |  |  |  |  |
| **DIC** | **0.90995** | **0.94602** | **0.95652** | **0.93492** | **0.85984** | **0.75294** | -0.07579 | **0.60394** | 0.15005 | **0** |  |  |  |  |
| **DIM** | **0.88986** | **0.91391** | **0.92763** | **0.8956** | **0.84656** | **0.75178** | -0.02914 | **0.59559** | 0.15018 | -0.11796 | **0** |  |  |  |
| **SGO** | **0.9703** | **1** | **1** | **1** | **0.93001** | **0.86799** | 0.14894 | **0.73231** | **0.25487** | 0.09091 | 0.02946 | **0** |  |  |
| **CIP** | **0.83454** | **0.85132** | **0.87422** | **0.82301** | **0.79267** | **0.6958** | **0.74785** | **0.82954** | **0.74843** | **0.8348** | **0.83193** | **0.90337** | **0** |  |
| **OUR** | **0.89449** | **0.91292** | **0.92573** | **0.89773** | **0.86103** | **0.78583** | **0.81997** | **0.87776** | **0.81378** | **0.89335** | **0.88378** | **0.94245** | **0.30238** | **0** |

JAC: Jacobina - BA; MKA: Miguel Calmon - BA; MCH: Morro do Chapéu - BA; MUC: Mucugê - BA; RCO: Rio de Contas - BA; ABA: Abaíra - BA; LIC: Licínio de Almeida - BA; GMO: Grão Mogol - MG; DIB: Diamantina (Biribiri) - MG; DIC: Diamantina (Cons. Mata) - MG; DIM: Diamantina (Milho Verde) - MG; SGO: São Gonçalo do Rio Preto - MG; CIP: Conceição do Mato Dentro - MG; OUR: Ouro Branco – MG.

Table S5.3 - Pairwise F_ST_ among 12 populations of Vriesea oligantha based on nrDNA microsatellites. All significant values are in bold.

|  | **JAC** | **MCH** | **MUC** | **ABA** | **RCO** | **LIC** | **GMO** | **DIB** | **DIC** | **SGO** | **CIP** | **OUR** |
| --- | --- | --- | --- | --- | --- | --- | --- | --- | --- | --- | --- | --- |
| **JAC** | **0** |  |  |  |  |  |  |  |  |  |  |  |
| **MCH** | **0.60448** | **0** |  |  |  |  |  |  |  |  |  |  |
| **MUC** | **0.27558** | **0.55063** | **0** |  |  |  |  |  |  |  |  |  |
| **ABA** | **0.40826** | **0.58123** | **0.44628** | **0** |  |  |  |  |  |  |  |  |
| **RCO** | **0.43111** | **0.68538** | **0.45668** | 0.03048 | **0** |  |  |  |  |  |  |  |
| **LIC** | **0.35194** | **0.73545** | **0.27868** | **0.5275** | **0.58596** | **0** |  |  |  |  |  |  |
| **GMO** | **0.4258** | **0.71828** | **0.44233** | **0.4974** | **0.54094** | **0.27769** | **0** |  |  |  |  |  |
| **DIB** | **0.28499** | **0.60217** | **0.28804** | **0.37852** | **0.40288** | **0.24587** | **0.19927** | **0** |  |  |  |  |
| **DIC** | **0.2765** | **0.63076** | **0.28076** | **0.39132** | **0.42489** | **0.30363** | **0.297** | 0.00813 | **0** |  |  |  |
| **SGO** | **0.30511** | **0.62592** | **0.29139** | **0.42439** | **0.45532** | **0.258** | **0.26111** | 0.01276 | 0.00786 | **0** |  |  |
| **CIP** | **0.40719** | **0.70189** | **0.41326** | **0.47976** | **0.52067** | **0.44173** | **0.29706** | **0.24608** | **0.2926** | **0.29227** | **0** |  |
| **OUR** | **0.51005** | **0.79466** | **0.48566** | **0.57567** | **0.63612** | **0.58351** | **0.39143** | **0.34761** | **0.42841** | **0.38705** | **0.07238** | **0** |

JAC: Jacobina - BA; MCH: Morro do Chapéu - BA; MUC: Mucugê - BA; RCO: Rio de Contas - BA; ABA: Abaíra - BA; LIC: Licínio de Almeida - BA; GMO: Grão Mogol - MG; DIB: Diamantina (Biribiri) - MG; DIC: Diamantina (Cons. Mata) - MG; SGO: São Gonçalo do Rio Preto - MG; CIP: Conceição do Mato Dentro - MG; OUR: Ouro Branco – MG.

Table S5.4 - Contemporary migration rates estimated in BAYESASS among 12 populations of *Vriesea oligantha*. Estimated migration rates > 0.10 are displayed in red.

| From/To | JAC | MCH | MUC | RCO | ABA | LIC | GMO | DIB | DIC | SGO | CIP | OUR |
| --- | --- | --- | --- | --- | --- | --- | --- | --- | --- | --- | --- | --- |
| JAC | 0.8666 | 0.0107 | 0.0112 | 0.0079 | 0.0107 | 0.0196 | 0.0104 | 0.0104 | 0.0112 | 0.0124 | 0.0095 | 0.009 |
| MCH | 0.0119 | 0.8819 | 0.0106 | 0.0081 | 0.0107 | 0.0196 | 0.0104 | 0.0104 | 0.0112 | 0.0124 | 0.0096 | 0.009 |
| MUC | 0.0124 | 0.0107 | 0.8777 | 0.008 | 0.0107 | 0.0196 | 0.0105 | 0.0104 | 0.0111 | 0.0124 | 0.0096 | 0.009 |
| RCO | 0.0119 | 0.0107 | 0.0107 | 0.9125 | 0.2153 | 0.0196 | 0.0104 | 0.0105 | 0.0111 | 0.0125 | 0.0095 | 0.009 |
| ABA | 0.0122 | 0.0108 | 0.011 | 0.0079 | 0.6774 | 0.0196 | 0.0104 | 0.0104 | 0.0111 | 0.0124 | 0.0095 | 0.009 |
| LIC | 0.0122 | 0.0107 | 0.0108 | 0.0079 | 0.0107 | 0.6863 | 0.0104 | 0.0104 | 0.0112 | 0.0124 | 0.0096 | 0.009 |
| GMO | 0.0119 | 0.0108 | 0.0123 | 0.008 | 0.0107 | 0.1176 | 0.8854 | 0.0127 | 0.0112 | 0.0136 | 0.0095 | 0.0089 |
| DIB | 0.0123 | 0.0108 | 0.011 | 0.008 | 0.0107 | 0.0196 | 0.0105 | 0.8709 | 0.2107 | 0.1954 | 0.0095 | 0.009 |
| DIC | 0.0122 | 0.0107 | 0.0109 | 0.0079 | 0.0107 | 0.0196 | 0.0105 | 0.0104 | 0.6778 | 0.0124 | 0.0095 | 0.009 |
| SGO | 0.0122 | 0.0107 | 0.011 | 0.0079 | 0.0107 | 0.0196 | 0.0104 | 0.0105 | 0.0111 | 0.6791 | 0.0096 | 0.009 |
| CIP | 0.0122 | 0.0108 | 0.011 | 0.008 | 0.0107 | 0.0196 | 0.0104 | 0.0105 | 0.0111 | 0.0124 | 0.6762 | 0.0091 |
| OUR | 0.0119 | 0.0107 | 0.0117 | 0.0079 | 0.0108 | 0.0196 | 0.0104 | 0.0227 | 0.0111 | 0.0126 | 0.2283 | 0.901 |

JAC: Jacobina - BA; MCH: Morro do Chapéu - BA; MUC: Mucugê - BA; RCO: Rio de Contas - BA; ABA: Abaíra - BA; LIC: Licínio de Almeida - BA; GMO: Grão Mogol - MG; DIB: Diamantina (Biribiri) - MG; DIC: Diamantina (Cons. Mata) - MG; SGO: São Gonçalo do Rio Preto - MG; CIP: Conceição do Mato Dentro - MG; OUR: Ouro Branco – MG

Table S5.5 -Posterior distribution values for the Neighbor groups only, estimated on Migrate.

| **Parameter** | **2.5%** | **25%** | **Mode** | **75%** | **97.50%** | **Median** | **Mean** |
| --- | --- | --- | --- | --- | --- | --- | --- |
| Θ_1_ | 0.800 | 1.400 | 1.760 | 2.066 | 2.466 | 1.766 | 1.681 |
| Θ_2_ | 2.000 | 2.333 | 2.633 | 2.933 | 3.666 | 2.766 | 2.511 |
| Θ_3_ | 4.200 | 4.200 | 4.300 | 4.733 | 5.533 | 4.566 | 4.370 |
| Θ_4_ | 0.800 | 1.400 | 1.766 | 2.066 | 2.533 | 1.766 | 1.707 |
| M_2->1_ | 1.200 | 1.640 | 1.823 | 1.960 | 2.160 | 1.763 | 1.733 |
| M_1->2_ | 0.020 | 0.120 | 0.190 | 0.247 | 0.373 | 0.203 | 0.202 |
| M_3->2_ | 0.287 | 0.427 | 0.503 | 0.580 | 0.713 | 0.510 | 0.506 |
| M_2->3_ | 0 | 0.113 | 0.170 | 0.173 | 0.173 | 0.137 | 0.135 |
| M_4->3_ | 0.073 | 0.187 | 0.257 | 0.327 | 0.473 | 0.277 | 0.273 |
| M_3->4_ | 0.380 | 0.540 | 0.623 | 0.760 | 0.967 | 0.677 | 0.676 |

Θ – Effective population size; M_A -> B_ – Migration from group A to B. Group numbers are 1. Northern Espinhaço (JAC and MKA); 2. Chapada Diamantina (MCH, MUC, RCO and ABA); 3. Diamantina Plateau (LIC, GMO, DIB, DIC, DIM and SGO); 4. Southern Espinhaço (CIP and OUR). JAC: Jacobina - BA; MCH: Morro do Chapéu - BA; MUC: Mucugê - BA; RCO: Rio de Contas - BA; ABA: Abaíra - BA; LIC: Licínio de Almeida - BA; GMO: Grão Mogol - MG; DIB: Diamantina (Biribiri) - MG; DIC: Diamantina (Cons. Mata) - MG; SGO: São Gonçalo do Rio Preto - MG; CIP: Conceição do Mato Dentro - MG; OUR: Ouro Branco – MG.

Table S5.6 – Probability of the data of the two tested models (marginal likelihood) estimated on MIGRATE.

| **Model** | **RTS** | **BAS** | **HM** | **LBF** | **MPP** |
| --- | --- | --- | --- | --- | --- |
| Panmictic | -472245 | -78485.9 | -1738.12 | 0 | 0 |
| Neighboors only | -428574 | -71604.3 | -1717.89 | -6881.56 | 1.00 |

RTS - Raw thermodynamic score; BAS - Bezier approximation score; HM – Harmonic mean; LBF – Log Bayes Factor; MPP – Model posterior probability

Table S5.7 – Probability of population reduction, estimated with BOTTLENECK 1.2.02, using all loci fitting the two phases model (TPM).

| **Population** | **Wilcoxon test (TPM)** |
| --- | --- |
| JAC | 0.15039 |
| MCH | 0.00098 |
| MUC | 0.02441 |
| RCO | 0.21289 |
| ABA | 0.27344 |
| LIC | 0.15625 |
| GMO | 0.41016 |
| DIB | 0.28516 |
| DIC | 0.15039 |
| SGO | 0.08203 |
| CIP | 0.32617 |
| OUR | 0.00195 |

JAC: Jacobina - BA; MCH: Morro do Chapéu - BA; MUC: Mucugê - BA; RCO: Rio de Contas - BA; ABA: Abaíra - BA; LIC: Licínio de Almeida - BA; GMO: Grão Mogol - MG; DIB: Diamantina (Biribiri) - MG; DIC: Diamantina (Cons. Mata) - MG; SGO: São Gonçalo do Rio Preto - MG; CIP: Conceição do Mato Dentro - MG; OUR: Ouro Branco – MG
